# Examining NF-κB genomic interactions by ChIP-seq and CUT&Tag

**DOI:** 10.1101/2024.08.11.607521

**Authors:** Allison E. Daly, Allison Schiffman, Alexander Hoffmann, Stephen T. Smale

## Abstract

An understanding of the mechanisms and logic by which transcription factors coordinate gene regulation requires delineation of their genomic interactions at a genome-wide scale. Chromatin immunoprecipitation-sequencing (ChIP-seq) and more recent techniques, including CUT&Tag, typically reveal thousands of genomic interactions by transcription factors, but without insight into their functional roles. Due to cost and time considerations, optimization of ChIP experimental conditions is typically carried out only with representative interaction sites rather than through genome-wide analyses. Here, we describe insights gained from the titration of two chemical crosslinking reagents in genome-wide ChIP-seq experiments examining two members of the NF-κB family of transcription factors: RelA and c-Rel. We also describe a comparison of ChIP-seq and CUT&Tag. Our results highlight the large impact of ChIP-seq experimental conditions on the number of interactions detected, on the enrichment of consensus and non-consensus DNA motifs for the factor, and on the frequency with which the genomic interactions detected are located near potential target genes. We also found considerable consistency between ChIP-seq and CUT&Tag results, but with a substantial fraction of genomic interactions detected with only one of the two techniques. Together, the results demonstrate the dramatic impact of experimental conditions on the results obtained in a genome-wide analysis of transcription factor binding, highlighting the need for further scrutiny of the functional significance of these condition-dependent differences.

## Background

The binding of hundreds of proteins to genomic DNA and chromatin helps regulate differential transcription and several other molecular processes, including DNA replication, recombination, repair, and transposition. For many years, the specific genomic locations associated with proteins of interest could be detected only using in vitro or non-physiological in vivo approaches. However, beginning in the mid-1980s, the development of methods that allow researchers to monitor the genomic locations of protein interactions in physiological settings and at a genome-wide scale revolutionized the molecular biology field.

In the method used most frequently, known as chromatin immunoprecipitation (ChIP), the chromatin is first isolated from cells and fragmented by sonication or nuclease digestion, followed by immunoprecipitation using antibodies that recognize the protein (or post-translationally modified protein) of interest [1]. Immunoprecipitation enriches for protein-associated DNA fragments in the immunoprecipitation pellet, with the DNA fragments originally detected by hybridization [2, 3], and later by polymerase chain reaction (PCR) [4–7]. To detect protein-associated DNA fragments at a genome-wide scale, fragment cloning and DNA microarrays were initially employed [8–14] but were subsequently replaced by high-throughput sequencing in a method known as ChIP-seq [15, 16].

Because most sequence-specific DNA-binding proteins examined by ChIP do not bind DNA with sufficient stability to remain bound during chromatin isolation, fragmentation, and immunoprecipitation, proteins are typically covalently crosslinked to the DNA and to other nearby chromatin-associated proteins. The first ChIP experiments relied on ultraviolet irradiation to catalyze crosslinking [2, 3] but formaldehyde soon emerged as an attractive crosslinking agent due to its ability to covalently link the amino or imino groups of DNA bases to amino acids of closely associated proteins, most commonly the ε-amine of lysine [17, 18].

Nevertheless, for many DNA-binding proteins, formaldehyde alone does not allow protein-DNA crosslinking with sufficient efficiency for robust results, leading to the frequent addition of a second crosslinking reagent in ChIP protocols [19–22]. Disuccinimidyl glutarate (DSG, Pierce), a membrane-permeable homo-bifunctional N-hydroxysuccinimide ester crosslinker, is one such reagent. The amine-reactive esters of DSG catalyze protein-protein crosslinking and can enhance the efficiency of ChIP by linking a protein of interest that may not efficiently crosslink to DNA itself to nearby proteins that may be crosslinked to DNA more efficiently [20–22].

Although many laboratories have obtained important insights from ChIP and ChIP-seq experiments examining transcriptional regulators, the results remain challenging to interpret. One major challenge is that ChIP-seq experiments typically reveal interactions with hundreds or thousands of genomic locations. Several studies have suggested that a substantial fraction of the interactions detected in ChIP experiments may lack anticipated functional roles [23–26]. These interactions may instead correspond to low-specificity or low-affinity events as factors scan the genome for their functional interaction sites, or they may reflect broader roles of the factors in chromatin organization that remain to be elucidated. The fraction of interaction sites detected by ChIP-seq that are functionally significant remains unknown for all transcriptional regulators.

In an effort to circumvent the limitations of ChIP-seq and its frequent reliance on chemical crosslinking, a variety of alternative methods for examining protein-DNA interactions at a genome-wide scale have been developed (e.g. DamID [27]). Most recently, the CUT&Run and CUT&Tag methods have shown great promise [28, 29]. In CUT&Tag, primary antibodies to the protein of interest are first incubated with isolated nuclei, followed by incubation with secondary antibodies and Protein A/G fused to the Tn5 transposase, which interacts with the secondary antibody to cleave the DNA near the protein of interest. Transposase cleavage releases DNA fragments for library preparation and sequencing.

When initiating a ChIP-seq experiment, an important first step is to perform pilot experiments to optimize the procedure, often including a titration of crosslinker concentrations. These optimization steps are typically performed with representative DNA fragments that are already known to interact with the protein of interest, along with negative controls. Crosslinking titrations rarely involve full ChIP-seq experiments, often due to cost and time considerations. We are unaware of published reports describing the impact of varying crosslinker concentrations on ChIP-seq results.

The nuclear factor κB (NF-κB) family of transcription factors consists of five family members in most vertebrate species [30, 31]. Each family member contains a conserved Rel homology region (RHR) that supports sequence-specific DNA binding and the formation of a variety of homodimers and heterodimers [31]. NF-κB dimers are thought to contribute to the transcriptional activation of a large number of genes by binding DNA recognition motifs within gene promoters and enhancers. Most dimers are activated in response to environmental stimuli, including microbial pathogens, radiation, and other inflammatory cues [31]. Although NF-κB dimers are among the most widely studied transcription factors, much remains to be learned about the mechanisms by which they coordinate transcriptional responses.

Genome-wide ChIP-chip and ChIP-seq experiments with NF-κB family members have been reported in mouse and human cells [24, 32–35], revealing interactions at promoters and enhancers containing and lacking NF-κB consensus recognition sequences. However, as with other transcriptional factors, thousands of genomic interactions are typically detected, many at locations lacking consensus motifs, with little knowledge of the functional roles of the vast majority of these interactions. To increase our understanding of NF-κB regulatory mechanisms, it is important to understand how the genomic interaction landscape is influenced by crosslinking conditions and the genomic interaction method. This understanding was critical for defining preferred genome profiling conditions in our lab for examining the specific functions and mechanisms of action of distinct NF-κB subunits and dimeric species [36, 37]. Here, we describe the results and insights obtained when varying chemical crosslinker concentrations for ChIP-seq experiments examining the RelA and c-Rel members of the NF-κB family, and when comparing ChIP-seq to CUT&Tag, a method that does not involve chemical crosslinking.

## Methods

### Cell culture

Bone marrow-derived macrophages (BMDMs) were prepared as described [38] from C57BL/6 male mice 8-12 weeks of age. Following extraction of the bone marrow, cells were incubated for 4 days with 10% CMG-conditioned media to begin differentiation into macrophages. On day 4, cells were scraped and plated at 5x10^6^ cells per 15 cm plate in fresh media containing 10% CMG. On day 6, cells were stimulated with 100 ng/mL lipid A (Sigma) for 60 mins. HoxB4-transduced myeloid progenitor-derived macrophages (hMPDMs) were differentiated from HoxB4-transduced myeloid progenitors as described [39–41] with medium containing 10% ES FBS, 1% penicillin/streptomycin, 1% L-glutamine, 30% L929 cell supernatant, and 2-mercaptoethanol (1:1000). 200,000 cells were plated in 2 ml of differentiation media [41] for each time point. On day 7 of differentiation, hMPDMs were stimulated with 100 ng/mL LPS (Sigma Aldrich).

### RNA-seq

Chromatin-associated RNA-seq data are from Tong et al. [34] (GEO accession GSE67357). Reads were aligned with Hisat2 to the NCBI37/mm9 genome. RPKM values were calculated for each sample by dividing the total reads for one gene by the length of the gene in kbps and the total reads per sample. Gene induction was calculated by averaging the RPKM values for three biological replicates at the 0 h and 1 h lipid A stimulation time points and taking the ratio of 1 h average RPKM versus the 0 h average RPKM.

### ChIP-seq

ChIP-seq was performed as previously described [33, 42] with anti-RelA (Cell Signaling, 8242) or anti-c-Rel antibody (Cell Signaling, 67489). The c-Rel antibody was validated by ChIP-seq performed with *Rel^-/-^* BMDMs (data not shown). Approximately 20 million BMDMs were used per sample. After cross-linking with DSG (0-4 mM) and formaldehyde (0-2%), cells were sonicated on a Covaris M220-focused ultrasonicator. The proper distribution of chromatin was checked using DNA electrophoresis to ensure DNA fragmentation was between 200-500bps.

ChIP-seq libraries were prepared using KAPA HyperPrep Kits (Roche) and barcode indices from NextFlex (Perkin Elmer). Following sequencing and demultiplexing, reads were aligned using Hisat2 to the NCIB/mm9 mouse genome. Peak calling was performed with Homer software, using input samples to find peak enrichment with a p-value < 0.01 (Heinz et al., 2010). To compare peaks across multiple samples, a master probe was generated with BEDTools [43]. Then RPKMs were generated using raw reads from SeqMonk (Babraham Bioinformatics) normalized to the size of the peak (in kbps) and the depth of sample sequencing (in million reads).

### CUT&Tag

CUT&Tag [29] was performed on 100,000 cells with CUTANA CUT&Tag reagents following manufacturer’s recommendations (EpiCypher CUTANA Direct-to-PCR CUT&Tag Protocol v1.7), with an anti-RelA antibody (Santa Cruz Biotech sc-109). Library preparation was performed with 0.4 μM each of universal i5 primer and barcoded i7 primers in NEBNext PCR master-mix for 16 cycles. The final DNA library was isolated with 1.3x AMPure beads. Sequencing was performed on a HiSeq 3000 to generate 50-bp single-end reads with about 10 million reads per sample. Reads were analyzed as described above for ChIP-seq reads. All high-throughput sequencing data for both ChIP-seq and CUT&Tag can be found under GEO accession GSE249834.

## Results

### Impact of varying the concentration of the protein-protein crosslinker DSG

To examine the impact of chemical crosslinking conditions on NF-κB ChIP-seq results, we performed ChIP-seq with different concentrations of both formaldehyde and DSG (see above), with a focus on two NF-κB family members that possess similar DNA-binding specificities, RelA and c-Rel [44]. Although varying the time and temperature of crosslinking was found to also influence ChIP-seq results (data not shown), we found the largest impacts when varying the concentrations of the chemical crosslinkers.

To first examine the impact of DSG concentration on ChIP-seq results, ChIP-seq for RelA and c-Rel was performed with 0, 1, 2, and 4 mM DSG, which spans the range of 0-5 mM typically recommended in ChIP-seq protocols. In all experiments, 1% formaldehyde was included. The experiments were performed with mouse BMDMs stimulated for 0 or 1 h with lipid A, a microbial product that engages Toll-like receptor 4 (TLR4) to potently activate NF-κB dimers via their nuclear translocation [34].

The ChIP-seq results reveal a dramatic increase in the number of statistically called peaks for both c-Rel and RelA 1-h post-stimulation with increasing DSG concentration (Fig. 1A). For example, 2,971 and 67,961 RelA peaks (p-adj < 0.01; peak score >19) are observed with 0 mM and 4 mM DSG, respectively. As expected, fewer than 100 called peaks are observed for both RelA and c-Rel in unstimulated cells (data not shown). For each DSG concentration examined, the peaks were separated into two groups: 1. peaks detected both with the specified DSG concentration and with any lower concentration of DSG (Fig. 1A, “previous peaks”) and 2. peaks that are called only with the specified DSG concentration (Fig. 1A, “new peaks”). With each DSG concentration, almost all peaks observed with lower DSG concentrations are detected, as expected, in addition to a large number of new peaks.

**Fig. 1.**
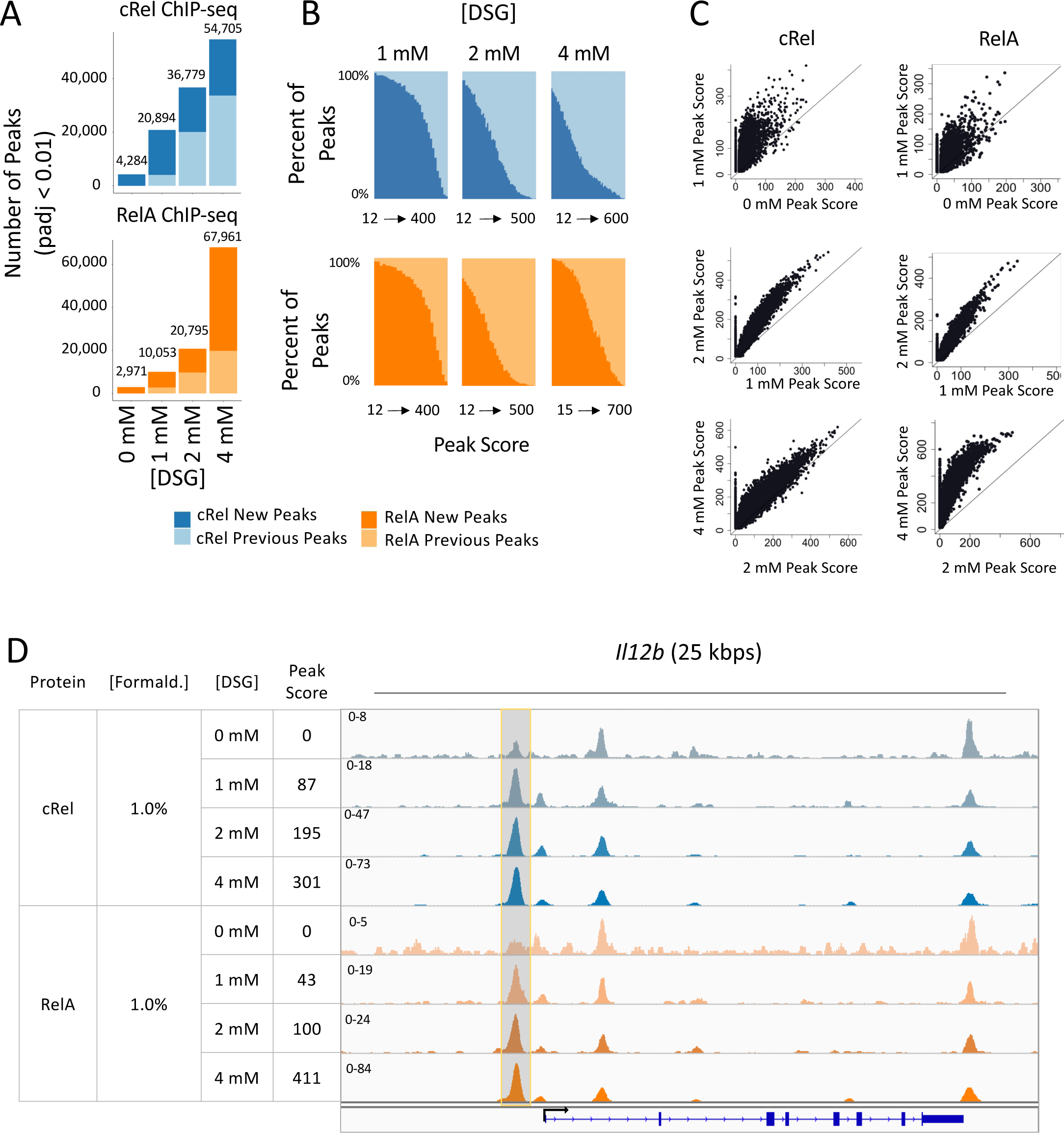
Impact of DSG concentration on RelA and c-Rel ChIP-seq results. BMDMs stimulated with lipid A for 1 hr were crosslinked with 0, 1, 2, or 4 mM DSG and 1% formaldehyde and examined by ChIP-seq. The results shown are from a single replicate of each DSG concentration. The findings are consistent with those of other experiments performed with variable crosslinker concentrations (data not shown). (**A**) The total number of peaks (p-adj <0.01; peak score > 19) for each condition is plotted. For each DSG concentration, peaks are further grouped into two categories: peaks called only at the specified DSG concentration (darker color, new peaks) or peaks called with both the specified concentration and any lower concentration of DSG (lighter color, previous peaks). Analyses for RelA and c-Rel peaks are colored in orange and blue, respectively. (**B**) All peaks are ranked based on peak score and separated into bins with 1,000 peaks per bin. The percentages of new peaks (darker color) and previous peaks (lighter color) are plotted. (**C**) The scatter plots show the peak scores for each called peak at each concentration of DSG. The plots show all peaks that are bound in either condition with a peak score > 0. (**D**) The Integrative Genomics Viewer (IGV) tracks spanning the *Il12b* locus are shown for c-Rel and RelA ChIP-seq with 0, 1, 2, or 4 mM DSG. A representative peak that is not called at the 0 mM concentration for c-Rel and RelA but is called at higher concentrations is highlighted in grey.

A peak located ∼1,000 bps upstream of the transcription start site (TSS) of the *Il12b* gene provides an example of the above findings (Fig. 1D, highlighted peak). This peak is not detected as a statistically called peak with 0 mM DSG for c-Rel or RelA. However, with 1 mM DSG, peaks for both c-Rel and RelA are detected, with the peak score increasing at higher DSG concentrations. This peak would be referred to as a “new peak” at the 1 mM concentration and as a “previous peak” with 2 and 4 mM DSG (Fig. 1D).

To determine the relationship between the peak scores at new versus previous peaks, we ranked the peaks for each DSG concentration based on peak score. We then grouped the peaks into bins containing 1,000 peaks each and created bar graphs showing, for each bin (x-axis) the percentages of peaks that correspond to previous peaks and new peaks (y-axis). This analysis reveals that new peaks at higher concentrations of DSG are biased toward low peak scores (Fig. 1B). For example, at 2 mM DSG, new peaks account for 98% of peaks in the bin with the lowest peak scores and 0% of the peaks in the bin with the highest peak scores (Fig. 1B). This finding suggests that, while more peaks are observed with higher DSG concentrations, these peaks are often weak and their functional roles and relevance may therefore require even closer scrutiny.

Scatter plots provide additional insights into the impact of increasing DSG concentration (Fig. 1C). Most peaks detected with a given DSG concentration exhibit comparable increases in peak score at the next highest concentration, with the most variability observed when the 0 and 1 mM DSG concentrations are compared (i.e. a more diffuse diagonal is observed with the 0 versus 1 mM comparison than with the other comparisons). However, these scatter plots also reveal the large numbers of new peaks with the higher DSG concentrations that were not detected with the lower concentration. That is, each plot shows a clear vertical line of peaks with peak scores of 0 on the x-axis, with thousands of overlapping peaks within these vertical lines. Although the overall correlation coefficient between neighboring conditions is high (data not shown), the slope of the best-fit line reveals that, on average, the peak scores approximately double with each increased DSG concentration (data not shown). Thus, increasing the DSG concentration generally increases existing peak scores relatively proportionally, while also revealing a large number of new peaks, generally with low peak scores.

### NF-κB consensus motif enrichment with increasing DSG concentrations

To further explore the genomic interactions detected with different DSG concentrations, we examined the enrichment of canonical NF-κB motifs in the collection of called peaks at each DSG concentration. Beginning with an analysis of known motifs using the HOMER program [45], which includes three slightly different NF-κB consensus motifs, we observed the strongest enrichment of all three NF-κB motifs when examining peaks called at 0 mM DSG and new peaks at 1 mM DSG (Fig. 2A). Enrichment of these motifs declines substantially in new peaks observed with 2 mM and 4 mM DSG. (Note that the same number of peaks [1000] were compared in each group.) Peaks called at 0 mM DSG, but not at the other concentrations, also exhibit enrichment of POU and Stat motifs, for reasons that remain unknown. New peaks observed with 4 mM DSG exhibit enrichment of zinc finger motifs, again for unknown reasons (Fig. 2A).

**Fig. 2.**
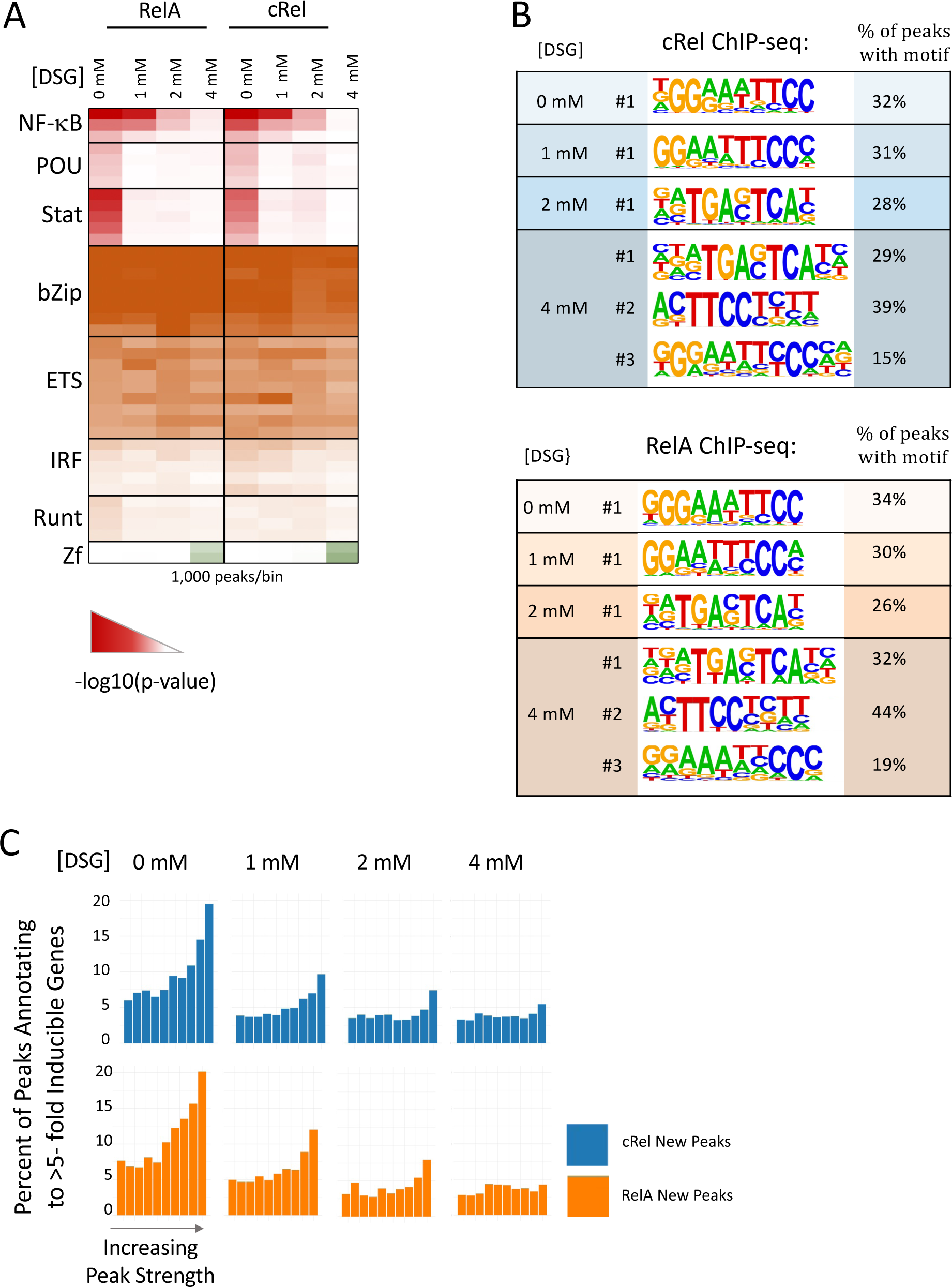
Impact of DSG concentration on motif enrichment and peak proximity to inducible genes. (**A**) Called peaks observed with 0 mM DSG and new peaks called when using 1, 2, and 4 mM DSG for both RelA and c-Rel ChIP-seq were examined. A random sample of 1,000 peaks from each group was used for known motif analysis using the HOMER program. The heat map corresponds to the –log10(p-value) and shows known motifs within eight different transcription factor families that exhibited the greatest enrichment in this analysis. Several rows are shown for each family, representing motifs within the HOMER program for different family members or dimeric species. (**B**) The peak groups described above were analyzed with HOMER de novo motif analysis software. The top motif in each analysis is shown for each DSG concentration and the top three motifs are shown for the 4 mM DSG samples, along with the percentage of peaks within each group containing a sequence matching the indicated motif. (**C**) Each peak within the group groups described above was annotated to its nearest gene. Peaks in each group were ranked based on peak score and grouped into 10 bins. The percentage of peaks in each bin that annotates to an inducible gene (induction > 5) based on nascent transcript RNA-seq analysis of BMDMs stimulated for 0 or 1 hr with lipid A are shown.

We also performed de novo motif analysis with new peaks observed at each DSG concentration (Fig. 2B). This analysis confirms strong enrichment of a consensus NF-κB motifs among 0 mM DSG peaks and new peaks observed with 1 mM DSG (Fig. 2B). (For the most abundant NF-κB dimer, a RelA:p50 heterodimer, the consensus motif defined biochemically is 5’-GGGRN(Y)YYCC-3’). However, at 2 and 4 mM DSG, a motif that resembles an NF-κB consensus motif is only ranked third among the enriched motifs and this enriched motif is much less rigid than the consensus NF-κB motif enriched with 0mM and 1mM DSG (Fig. 2B and data not shown). For example, note the increased flexibility of the GG dinucleotide within the first half-site in the third ranked motifs with 4mM DSG in comparison to the greater consistency of this dinucleotide in the motif enriched with 0mM and 1mM DSG. These results reveal that, although increasing DSG concentrations greatly increases the number of called peaks in both c-Rel and RelA ChIP-seq experiments, the newly detected peaks are strongly biased toward peaks with low peak scores (Fig. 1B) and they also coincide with motifs that exhibit greater divergence from an NF-κB consensus motif.

It may be noteworthy that the de novo motif exhibiting the greatest enrichment among newly called peaks with 2 and 4 mM DSG resembles consensus binding motifs for bZIP proteins. bZIP motifs are often enriched in regulatory regions induced by inflammatory stimuli [34, 35, 45, 46], raising the possibility that the new RelA and c-Rel ChIP-seq peaks observed with high DSG concentrations occur preferentially at regulatory regions that support inducible transcription. However, bZIP motifs are among the most enriched motifs at all constitutively open regulatory regions in macrophages [47]. This suggests that the bZIP enrichment at high DSG concentrations may instead be due to broad RelA and c-Rel crosslinking to open chromatin throughout the genome (see below).

Finally, the motif ranked second with 2 mM and 4 mM DSG resembles an NF-κB half-site and therefore, like the third-ranked motif, it could represent binding of NF-κB to lower affinity motifs. However, this motif also resembles a recognition motif for the transcription factor PU.1 that plays a role in macrophage development and is prevalent among macrophage enhancers.

### The proximity of RelA and c-Rel ChIP-seq peaks to inducible genes

To further address the characteristics of RelA and c-Rel ChIP-seq peaks observed at different DSG concentrations, we first divided all new peaks at each DSG concentration into ten bins based on peak score. We then annotated each peak in each bin to the nearest gene. Taking advantage of our previously reported nascent transcript RNA-seq data sets generated with unstimulated and lipid A-stimulated BMDMs [34], we calculated the percentage of peaks within each bin that annotate to a gene whose transcript levels were induced >5-fold following lipid A stimulation for 1 h (Fig. 2C).

This analysis reveals that the percentage of both RelA and c-Rel peaks annotating to strongly induced genes is much higher with 0 mM DSG than with new peaks observed at any of the higher DSG concentrations (Fig. 2C). In the 0 mM DSG bin containing peaks with the highest scores, approximately 20% of the peaks annotate to strongly induced genes. In new peaks detected with 1 mM DSG, 10-12% of peaks in the RelA and c-Rel bins with the highest peak scores annotate to induced genes, with only about 5% in the bins containing the strongest new peaks detected at 2 or 4 mM DSG. Notably, in the 0 mM DSG bins with the lowest peak scores, 7-8% of peaks annotate near induced genes, which is much larger than the 3-4% observed with new peaks observed with 4 mM DSG. The 3-4% observed with 4 mM DSG may represent a baseline (background) value.

Thus, if proximity to an inducible gene is viewed as a preliminary estimate of a functional role in inducible transcription, peaks called with 0 mM DSG have a higher probability of supporting this function regardless of peak score compared to new peaks called with 4 mM DSG. However, even with 0 mM DSG, the probability of a functional role appears to increase with increasing peak score. Notably, new peaks called with 1 mM DSG and to a lesser extent with 2 mM DSG also annotate near inducible genes with a higher prevalence than the apparent baseline level, but primarily in bins with the strongest peak scores.

It is also noteworthy that the new peaks observed with 4 mM DSG have a low probability of annotating near inducible genes despite the enrichment of bZIP motifs at these peaks (Fig. 2A and 2B). This finding strengthens the argument against the notion that new peaks detected with 4 mM DSG correspond to regulatory regions involved in inducible transcription (see above). Together, the results suggest that the large number of new peaks called with high DSG concentrations, especially 2 and 4 mM DSG, possess a relatively low probability of supporting inducible transcription. These peaks not only are generally weak and less likely to coincide with consensus NF-κB motifs, but they also are less likely to annotate to inducible genes than peaks observed with 0 mM and 1 mM DSG.

### Impact of varying the concentration of formaldehyde

Formaldehyde can catalyze both protein-protein and protein-DNA crosslinking but has emerged as a preferred crosslinking agent for ChIP-seq experiments due to the latter activity (Solomon et al. 1988). Because standard ChIP-seq protocols include 0.5% - 2.0% formaldehyde, we performed ChIP-seq experiments with 0.0%, 0.5%, 1.0%, and 2.0% formaldehyde in combination with 1 mM DSG for both c-Rel and RelA in BMDMs stimulated with lipid A for 0 or 1 h. We first assessed the overall number of statistically called peaks (p-adj < 0.01; peak score >19) in each condition (Fig. 3A). In contrast to the large number of peaks obtained in the absence of DSG, only 55 and 37 peaks are observed for c-Rel and RelA, respectively, in the absence of formaldehyde. With the addition of 0.5% formaldehyde, we observe a dramatic increase in the number of peaks, which continued to increase with higher formaldehyde concentrations (Fig. 3A).

**Fig. 3.**
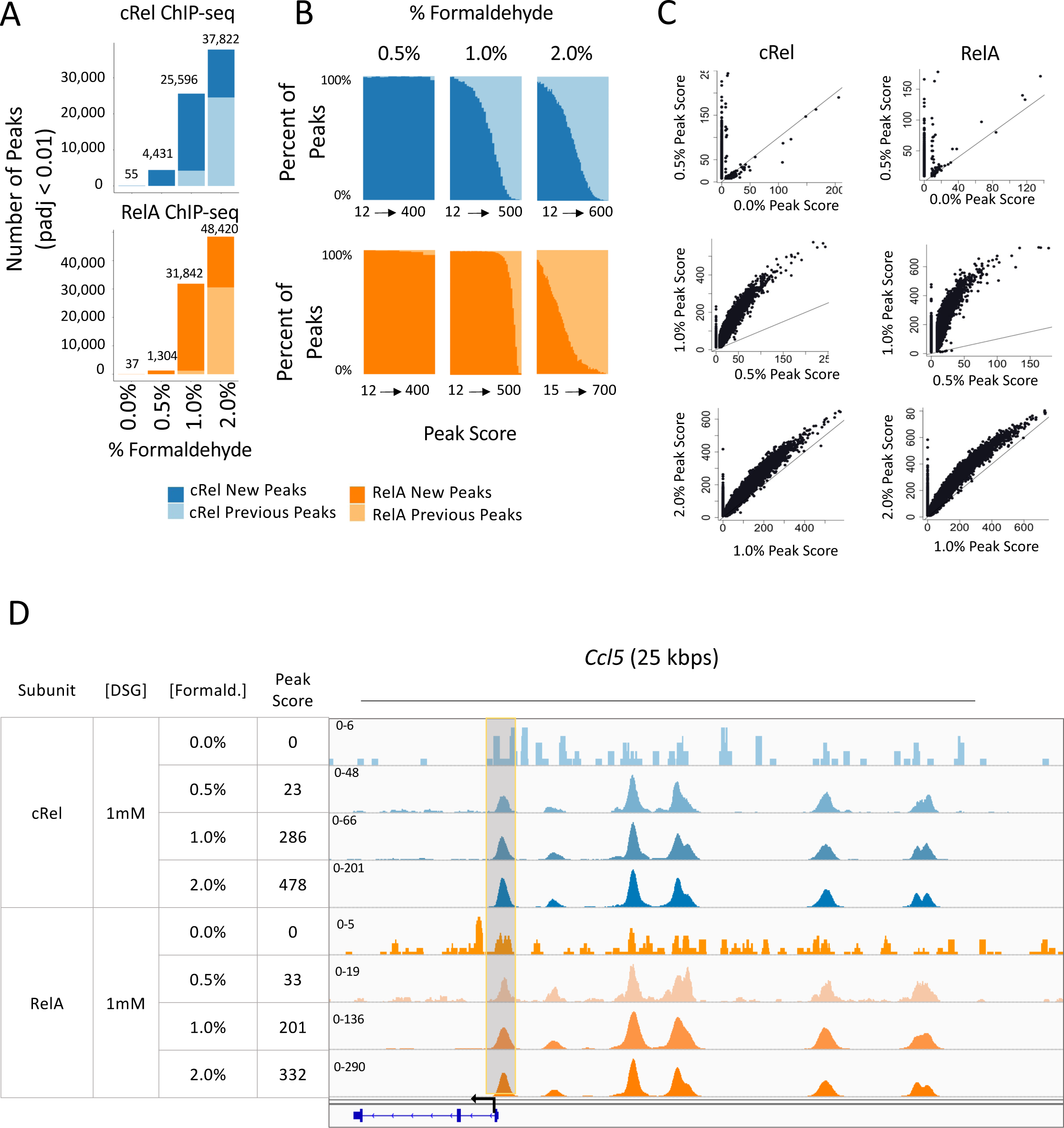
Impact of formaldehyde concentration on RelA and c-Rel ChIP-seq results. BMDMs stimulated with lipid A for 1 hr were crosslinked with 0, 0.5, 1.0, or 2.0% formaldehyde and 1 mM DSG and examined by ChIP-seq. The results shown are from a single replicate of each formaldehyde concentration. The findings are consistent with those of other experiments performed with variable crosslinker concentrations (data not shown). (**A**) The total number of peaks (p-adj <0.01; peak score > 19) for each condition is plotted. For each formaldehyde concentration, peaks are further grouped into two categories: peaks called only at the specified formaldehyde concentration (darker color, new peaks) or peaks called with both the specified concentration and any lower concentration of formaldehyde (lighter color, previous peaks). Analyses for RelA and c-Rel peaks are colored in orange and blue, respectively. (**B**) All peaks are ranked based on peak score and separated into bins with 1,000 peaks per bin. The percentages of new peaks (darker color) and previous peaks (lighter color) are plotted. (**C**) The scatter plots show the peak scores for each called peak at each concentration of formaldehyde. The plots show all peaks that are bound in either condition with a peak score > 0. (**D**) The IGV tracks spanning the *Ccl5* locus are shown for c-Rel and RelA ChIP-seq with each formaldehyde concentration. A representative peak that is not called at the 0% formaldehyde concentration for c-Rel and RelA but is called at higher concentrations is highlighted in grey.

Similar to the DSG analyses, increasing the formaldehyde concentration yields many of the same peaks observed with the lower formaldehyde concentrations, but also a large number of new peaks (Fig. 3A). Also similar to the DSG results, the new peaks with higher formaldehyde concentrations are generally weak (Fig. 3B). When using scatter plots to examine the impact of increased formaldehyde concentrations on peak scores with individual peaks, peaks observed with the lower formaldehyde concentration generally increase their peak score with the higher concentration, in addition to the appearance of the new peaks (Fig. 3C). This increase in the peak score for pre-existing peaks is most pronounced when the formaldehyde concentration was increased from 0.5% to 1%. The *Ccl5* promoter provides an example of a peak that first appears with 0.5% formaldehyde and continues to increase in peak score at 1% and 2% formaldehyde (Fig. 3D).

We also examined motif enrichment at new peaks with each formaldehyde concentration. The most significant enrichment of a consensus NF-κB motif is observed with new peaks called with 0.5% formaldehyde (Fig. 4A). This consensus motif is also enriched in new peaks called with 1% and 2% formaldehyde, but to a lesser extent. In a de novo motif analysis, a motif resembling an NF-κB consensus exhibits the greatest enrichment with the new peaks observed with 0.5% formaldehyde (Fig. 4B). However, motifs resembling an NF-κB consensus are more weakly enriched among new peaks observed with 1% and 2% formaldehyde, with the most enriched motif suggestive of bZIP family protein binding (Fig. 4B and data not shown). Given the very small number of peaks called with 0% formaldehyde, which are likely to represent background, it may be unsurprising that the most enriched motif had little resemblance to an NF-κB motif.

**Fig. 4.**
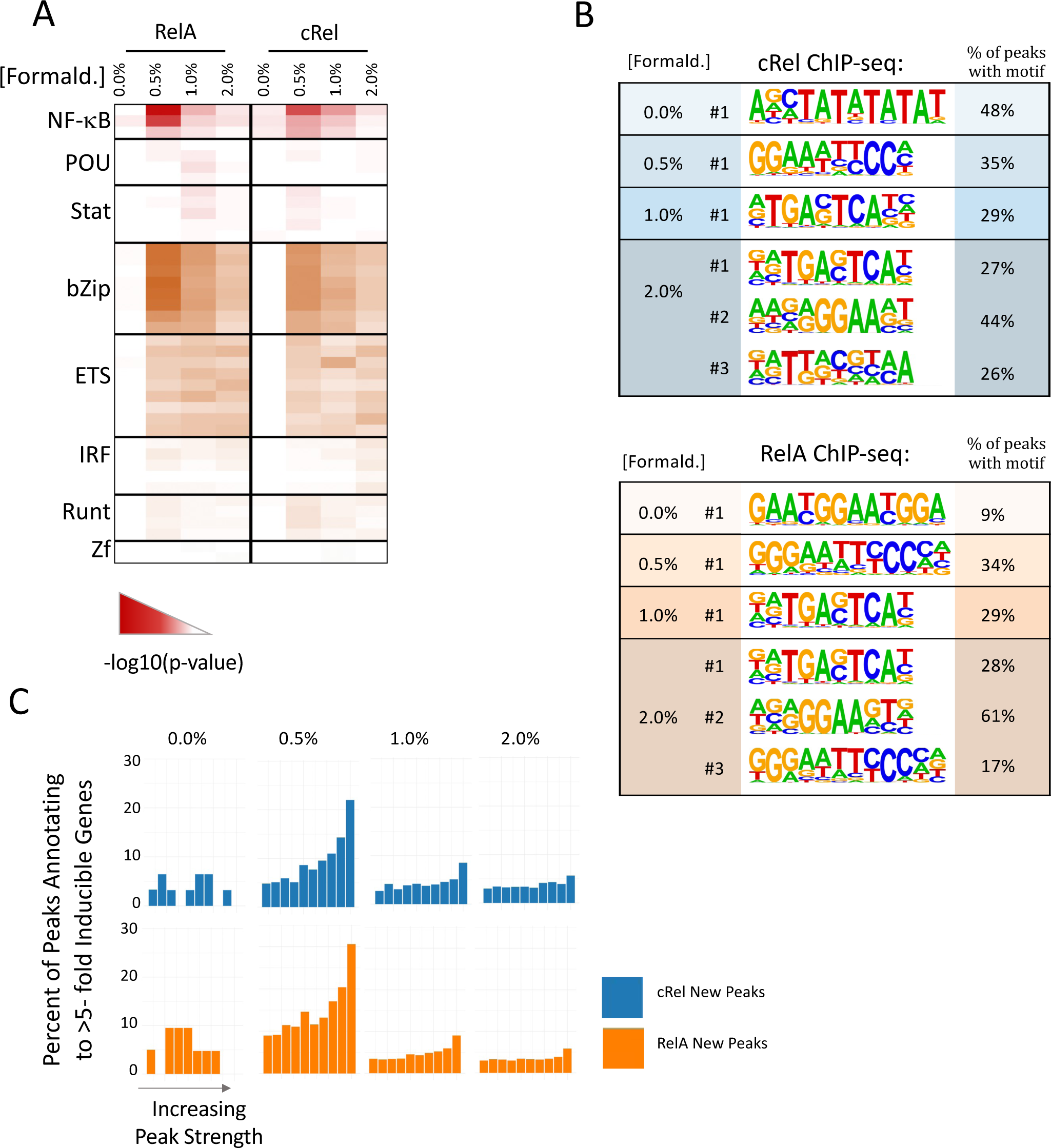
Impact of formaldehyde concentration on motif enrichment and peak proximity to inducible genes. (**A**) Called peaks observed with 0% formaldehyde and new peaks called when using 0.5, 1, and 2% formaldehyde for both RelA and c-Rel ChIP-seq were examined. A random sample of 1,000 peaks from each group was used for known motif analysis using the HOMER program. The heat map corresponds to the –log10(p-value) and shows known motifs within eight different transcription factor families that exhibited the greatest enrichment in this analysis. Several rows are shown for each family, representing motifs within the Homer program for different family members or dimeric species. (**B**) The peak groups described above were analyzed with HOMER de novo motif analysis software. The top motif in each analysis is shown for each DSG concentration and the top three motifs are shown for the 2% formaldehyde samples, along with the percentage of peaks within each group containing a sequence matching the indicated motif. (**C**) Each peak within the group groups described above was annotated to its nearest gene. Peaks in each group were ranked based on peak score and grouped into 10 bins. The percentage of peaks in each bin that annotates to an inducible gene (induction > 5) based on nascent transcript RNA-seq analysis of BMDMs stimulated for 0 or 1 hr with lipid A are shown.

Finally, new peaks called with each formaldehyde concentration were merged with nascent transcript RNA-seq data to determine the frequency with which they annotate to inducible genes. This analysis reveals that the newly called peaks observed with 0.5% formaldehyde exhibit a relatively high probability of annotating to induced genes in comparison to newly called peaks with 1% or 2% formaldehyde. Peaks in bins with the highest scores exhibit the highest probability of annotating to inducible genes. Together, these results suggest that, for both RelA and c-Rel, 0.5% formaldehyde appears to capture a large fraction of ChIP-seq peaks that have the highest probability of supporting inducible transcription, as measured by annotation to induced genes and by NF-κB motif enrichment and peak score. Notably, however, one major benefit of 1% formaldehyde is a large increase in peak score for those peaks initially detected with 0.5% formaldehyde (see Fig. 3C).

### ChIP-seq versus CUT&Tag comparison

Given the large impact of crosslinking conditions on the outcome of a ChIP-seq experiment, we compared ChIP-seq with CUT&Tag, a relatively new assay that does not use crosslinking [29]. For our comparison, CUT&Tag was performed with RelA hMPDMs, a mouse macrophage population that closely resembles BMDMs following their differentiation from HoxB4-transduced myeloid progenitors [39–41]. The hMPDMs were stimulated with lipid A for 15, 45, and 60 min, yielding 6,502, 6,616, and 4,809 CUT&TAG peaks, respectively.

The heat-maps in Fig. 5A display the comparison between the CUT&Tag peaks and the ChIP-seq peaks detected with different concentrations of DSG and formaldehyde. The heat-maps show the percent of peaks in the sample listed at the top that overlap with the sample listed to the left. For example, the 2 mM DSG ChIP-seq sample yielded 15,819 peaks (column 3). Most of these peaks are not detected with 0 mM DSG (column 3, row 1), thereby revealing limited overlap. A larger percentage of these 15,819 peaks overlaps with the 1 mM DSG peaks (column 3, row 2), with all 15,819 peaks overlapping with the same 2 mM DSG sample (column 3, row 3). Almost all of the 15,819 peaks are among the much larger number of peaks detected with 4 mM DSG (column 3, row 4).

**Fig. 5.**
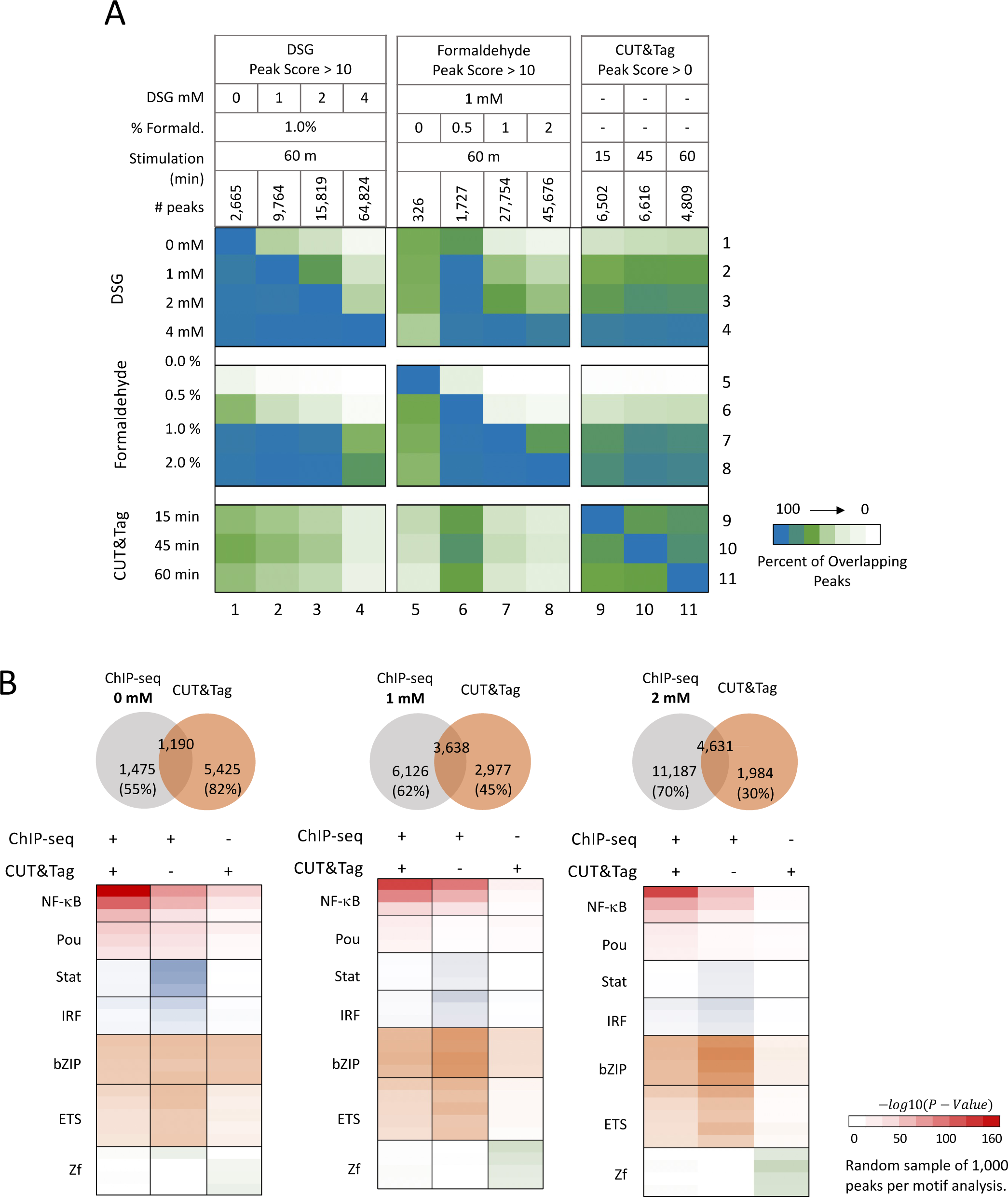
Comparing ChIP-seq to CUT&TAG. (**A**) The heat map shows the percent overlap of called peaks between each ChIP-seq sample and between each ChIP-seq sample and each CUT&Tag sample. The number of peaks that overlap between the two assays relative to the number of peaks in each sample (listed at the top of the chart) are shown as percentages in the heatmap (see Results section for further description). (**B**) The overlap between ChIP-seq peaks observed with three different DSG concentrations and the CUT&Tag peaks are shown as Venn diagrams (top). Known HOMER motif analysis was performed on peaks grouped into three categories: 1. peaks called in both the ChIP-seq (p-adj < 0.01) and CUT&Tag (p-adj < 0.01) experiments (with separate results shown for the three different DGS concentrations), 2. Peaks called only in the ChIP-seq experiments (p-adj <0.01), or 3. peaks called only in the CUT&Tag experiment (p-adj <0.01).

A close examination of the heat-maps reveals a number of insights. In particular, the three CUT&Tag datasets exhibit considerable overlap with each other (columns 9-11, rows 9-11). In addition, the three CUT&Tag datasets exhibit only moderate overlap with the ChIP-seq peaks observed with 0 mM DSG (columns 9-11, row 1), but increased overlap with the ChIP-seq peaks observed with 1 mM DSG (columns 9-11, row 2), with the overlap between these two samples comparable to the overlap observed between each of the three CUT&Tag samples (e.g. columns 10 and 11, row 9). (Note that the 1 mM DSG sample and the CUT&Tag samples also have comparable peak numbers.) A smaller percentage of the 9,764 peaks detected with 1 mM DSG overlaps with the three CUT&Tag datasets (column 2, rows 9-11) because of the larger number of peaks in the 1 mM DSG dataset. Overall, the overlap between the ChIP-seq datasets (e.g. with 1 mM DSG, 1% formaldehyde) and the CUT&Tag datasets is quite strong, confirming prior evidence of the similarity between the two methods [29].

Using peak score criteria (p-adj <0.01) to examine more closely the overlap between the ChIP-seq DSG titration data and the CUT&Tag data, we found that, while there is substantial overlap, there are also thousands of peaks that are observed only by either ChIP-seq or CUT&Tag (Fig. 5B, Venn diagrams at top). For example, in a comparison of 1 mM DSG ChIP-seq data with the CUT&Tag data, 3,638 peaks overlap, representing 38% and 55% of the ChIP-seq and CUT&Tag peaks, respectively (Fig. 5B, Venn diagrams top middle). This extensive overlap using two entirely different methodologies suggests that both techniques successfully capture NF-κB genomic binding sites. However, in this comparison, 62% and 45% of the ChIP-seq and CUT&Tag peaks, respectively, do not overlap with peaks obtained with the other technique.

To gain insight into the characteristics of the peaks detected by the two methods, we performed motif analysis with peaks detected in common by both methods, as well as peaks detected exclusively with one method (Fig. 5B, bottom). Peaks detected in common by both methods show strong enrichment of NF-κB motifs, regardless of whether the 0 mM, 1 mM, or 2 mM DSG data are used for the analysis, raising the possibility that interactions detected with both of these distinct methods have the highest probability of reflecting NF-κB bound to consensus sites (Fig. 5B, bottom). In contrast, weaker enrichment of NF-κB consensus motifs is observed at peaks detected with only one of the two techniques, with the weakest enrichment of NF-κB consensus motifs observed at peaks detected using only the CUT&Tag method. These results suggest that both methods generate peaks that do not represent the specific binding of NF-κB to consensus recognition motifs. Thus, such peaks are not dependent on chemical crosslinking.

## Discussion

In this study, we examined at a genome-wide scale how NF-κB protein:DNA interactions detected by ChIP-seq are impacted by chemical crosslinker concentrations and how NF-κB ChIP-seq and CUT&Tag results compare with each other. Although crosslinker titrations are frequently performed when developing a ChIP-seq assay for a new protein, these titrations are generally evaluated by PCR with only a small number of representative binding sites. We are unaware of other published reports describing the impact of crosslinker titrations at a genome-wide scale. We tested different concentrations of two chemical crosslinkers commonly used in ChIP-seq experiments and evaluated the impact on the overall ChIP-seq results and the specificity of interactions. The comparison between ChIP-seq and CUT&Tag results provided an opportunity not only to compare two entirely different methodologies, but also to compare a crosslinking-dependent method with a method that does not involve crosslinking.

Among the insights provided by these results are a demonstration of the difficult balance between optimizing ChIP-seq crosslinker concentrations for large numbers of peaks versus binding specificity and possible functional relevance. If proximity to inducible genes provides a rough predictor of the probability of functional relevance, crosslinking with formaldehyde alone appears to provide the highest confidence that called peaks may be functionally relevant. However, without the inclusion of a low concentration of DSG, a substantial number of interactions that are likely to be relevant would likely be missed. As the DSG concentration increases from 1 mM to 4 mM, the value of the increased number of peaks appears to decrease substantially, as higher and higher percentages of these peaks may not be functionally significant, or they may represent novel unappreciated activities of NF-κB. In fact, with 4 mM DSG, peaks are observed at over half of the regions of open chromatin detected in our lab by ATAC-seq [47] (data not shown).

Notably, in the absence of DSG, careful consideration of peak score also can influence the probability of functional relevance (once again, assuming that proximity to inducible genes provides a rough measure of the probability of functional relevance), as 2.5-3-fold more peaks with the highest peak scores are near inducible genes than observed with weak called peaks. It is already well-established that the detection of a protein-DNA interaction by ChIP-seq is insufficient for a conclusion that the interaction is functionally significant. This concept is strongly reinforced by our results.

For any given factor that interacts with DNA, there may be defined crosslinker concentrations that allow for the optimal capture of functionally important protein-DNA interactions, while minimizing to the greatest extent possible the capture of aberrant interactions. Further analyses are needed to understand the extent to which interactions captured by “optimal” crosslinker concentrations are functionally relevant, despite the strong enrichment of consensus motifs when using these conditions. Further analyses are also needed to determine whether the additional interactions captured with only higher crosslinker concentrations contribute to proper gene regulation. Despite the low prevalence of consensus recognition motifs and limited proximity to inducible genes, these interactions may somehow help support chromatin architecture within the nucleus.

The substantial overlap in interactions captured by ChIP-seq and CUT&Tag reinforces previous evidence that CUT&Tag is a highly valuable technique, given the method’s ease of use and suitability for use with small numbers of cells. However, both ChIP-seq and CUT&Tag captured large numbers of interactions that were not detected with the other method, and NF-κB consensus motifs exhibited much weaker enrichment at peaks captured with only one of the two methods. Although the significance of this finding is not known, one possible explanation is that each method has distinct susceptibilities to capturing interactions that are less likely to represent specific protein-DNA interactions mediated by direct physical interactions of RelA and c-Rel with DNA, in comparison to peaks that are reproducibly captured by both methods. Notably, those interactions detected by both of these two widely different methods may have the highest probability of representing specific binding events, and possibly functionally important binding events.

The use of chemical crosslinkers for ChIP-seq has long been known to provide susceptibility to background, due to the ability of a crosslinker to fix a highly transient interaction that may occur with little or no specificity. This concern is eliminated with CUT&Tag. However, the CUT&Tag method is susceptible to different potential challenges, including the potential for the transposase to preferentially cleave DNA at genomic sites without prior binding to the antibody bound to the protein of interest. Optimization of experimental conditions for CUT&Tag will help minimize these potential background cleavage events, but effective approaches will be needed to evaluate the results of optimization experiments. As done for the current analysis, evaluation of the optimal enrichment of consensus recognition motifs for a transcription factor of interest, or optimal enrichment of interactions near potential target genes, may be the preferred approach. However, this strategy will be of limited value for those transcription factors that may frequently carry out functional interactions with sites that diverge from their in vitro consensus recognition sequence, and it may lead investigators to overlook important unknown functions of factors that do not involve interactions with consensus motifs.

## Abbreviations

ChIP-seq: Chromatin immunoprecipitation sequencing
DSG: Disuccinimidyl glutarate
PCR: Polymerase chain reaction
NF-κB: Nuclear factor κB
BMDM: Bone marrow-derived macrophage
hMPDM: HoxB4-transduced myeloid progenitor-derived macrophage
TLR4: Toll-like receptor 4
TSS: Transcription start site
bZIP: Basic leucine zipper

## Acknowledgements

We thank Dr. Siavash Kurdistani for helpful comments on the manuscript.

## Author contributions

A.E.D. designed the experiments; acquired, analyzed, and interpreted the data; and prepared the manuscript. A.S. designed the experiments; and acquired, analyzed, and interpreted the data. A.H. designed the experiments; and interpreted the data. S.T.S. designed the experiments; interpreted the data; and prepared the manuscript.

## Funding

This work was supported by US National Institutes of Health grants R01GM086372, R01AI073868, R01CA127279, and P50AR063020 (to S.T.S.), and F31AI157267 and T32AI007323 (to A.E.D.). S.T.S. is a Senior Scientific Officer of the Howard Hughes Medical Institute.

## Data availability

High-throughput sequencing data for both ChIP-seq and CUT&Tag have been deposited and are publicly available at Gene Expression Omnibus accession number GSE249834.

## Declarations

### Competing interests

The authors declare no competing interests.

## Notes

### Competing Interest Statement

The authors have declared no competing interest.

